# Modeling human phenotypes in the nematode *C. elegans* during an undergraduate developmental biology course

**DOI:** 10.1101/2025.04.23.650323

**Authors:** Sireen Aburaide, Suad Ahmed, Mason Chamberlain, Yoskar Deleon, Ahmed Elewa, Allison Hookom, Evelyn Juen, Divine Katasi, Bobby Lee, Samantha Meyer, Luís Millan, Madeline Shaw, DJ Smith, Anthony Vassallo, Tashi Wangmo, Daniel Wolfson, Pader Xiong, Lian Yaeger, The Augsburg BIO474 Consortium

## Abstract

*Caenorhabditis elegans* is a widely used model organism in biomedical research due to its genetic tractability, short life cycle, and conservation of many developmental processes with humans. In this study, undergraduate students conducted nine independent experiments during a Developmental Biology course to model human phenotypes using *C. elegans*. Each group selected a human phenotype of interest, identified a gene associated with the phenotype, and then determined the orthologous or homologous gene in *C. elegans*. By obtaining mutants and designing phenotypic assays, students investigated the extent to which the worm models could recapitulate aspects of the human condition. This collective work highlights both the potential and limitations of *C. elegans* as a model for human phenotypic variation and disease and demonstrates the value of undergraduate inquiry as a catalyst for scientific engagement and research-based learning.

## Introduction

Beneath the superficial differences between species lurk the remains of common ancestry. Learning to trace and chase this yarn of evolution provides the student of nature with the skill to weave together disparate facts and compose our epic of descent with modification. Such learning is not attained by memorizing facts but by posing challenging questions, distilling principles, testing hypotheses and feeling the satisfaction of coherence post-confusion.

Therefore, integrating research into undergraduate education has emerged as a transformative approach to fostering scientific inquiry and critical thinking [1]. For instance, investigating the impact of course-based undergraduate research experiences (CUREs) revealed that students participating in full-semester CUREs reported greater gains in experimental design skills, career interests, and intentions to conduct future research compared to those in shorter module-based CUREs or traditional laboratory courses [2]. Similar initiatives collectively underscore a shift from traditional, didactic teaching to active, research-based education, where students are not passive learners but active contributors to scientific knowledge.

Here we report the experimental results of a research-based learning module as part of a 30-lecture course on Developmental Biology. Students independently selected phenotypes from the Human Phenotype Ontology database, selected associated human genes, and identified *C. elegans* orthologs or homologs for study [3,4]. *C. elegans* is widely recognized for its rapid life cycle, abundant experimental resources, and conservation of key developmental processes with humans, making it an ideal species for modeling human phenotypes and for undergraduate research [5-7]. Each group designed experiments using loss-of-function mutant strains to test whether the nematode could recapitulate aspects of the human condition. The results reflect a range of plausible and implausible models, revealing both opportunities and limits of modeling human phenotypes in *C. elegans* in an undergraduate setting.

## Methods

### Instruction in *C. elegans*

A fifteen week lab course was designed to complement thirty lectures in developmental biology (BIO474 Developmental Biology). During the first four weeks, students learned to pick *C. elegans* worms under a dissecting microscope and distinguish its different life stages. Students also learned how to mount worms on an agarose pad and observe fertilization and early embryonic cleavages under a compound microscope. Per worm tradition, students learned their worm lineage to Sydney Brenner. It was a particularly nice surprise to learn later on during the semester that Victor Ambros, a connection in their lineage, was a co-recipient of the 2024 Nobel Prize in Medicine or Physiology. Each student was responsible for their worm colony and maintained it by singling worms onto fresh seeded NGM plates and working with the resulting population the following week. A shared spreadsheet was used to log the overall state of each colony each week. <u>Between weeks 5 and 8</u>, students were grouped in pairs and each pair chose two human phenotypes of interest by browsing The Human Phenotype Ontology database. The sole criterion for choosing a human phenotype was curiosity. Students then were asked to choose a gene associated with the human phenotype of interest and present their choices to the class. Only afterwards were the students informed that they will now attempt to model the human phenotype via the associated gene in *C. elegans*. Students were encouraged to generate interesting hypotheses and if needed sufficiency abstract the phenotype in question to reach a conserved core developmental process. Finally, students sought *C. elegans* orthologs or homologs of their human gene of interest and identified suitable strains provided through the *Caenorhabditis* Genetics Center. By the end of this process, one of the two phenotypes originally chosen proved more promising and practical to pursue. In parallel to activities of this segment, students were also introduced to a host of *C. elegans* mutants and phenotypes to prepare them for work with non-wildtype strains. <u>Between weeks 9 and 15</u> and while waiting for strains to arrive, students designed experiments to test the extent to which the *C. elegans* strain of choice could model the human phenotype of interest. This iterative process came hand in hand with consulting course lectures, scientific literature via Pubmed [8] and the databases UniProt [9], AlphaFold [10,11] and WormBase [4] to learn more about the phenotype of interest, its associated diseases, the function of its associated gene and the documented phenotypes of the *C. elegans* homologs. The swift delivery of worm strains from CGC allowed each student group at least three weeks to troubleshoot and optimize their protocols and obtain data to evaluate their hypotheses.

### Strains

The following *C. elegans* strains used in this study were obtained from the *Caenorhabditis*

Genetics Center (University of Minnesota):

N2

CB1 *dpy-1(e1)* III. (Dpy)

CB1009 *unc-54(e1009)* I. (Unc - paralyzed)

CB1026 *lin-1(e1026)* IV. (Muv)

CB1138 *him-6(e1104)* IV. (Him)

CB1179 *unc-22(e1179)* IV. (Unc - twitcher)

CB1309 *lin-2(e1309)* X. (Vul)

CB1393 *daf-8(e1393)* I. (Daf)

CB1515 *mec-10(e1515)* X. (Mec)

CB1598 *unc-1(e1598)* X. (Unc - coiler)

CB91 *rol-1(e91)* II. (Rol)

RB1565 *ZK1073*.*1(ok1899)* X. Not outcrossed.

VC1224 *polg-1(ok1548)*/mT1 II; +/mT1 [*dpy-10(e128)*] II. *polg-1(ok1548)* outcrossed once.

JD21 *cca-1(ad1650)* X. Outcrossed 7 times.

GG25 *let-2(g25)* X. Outcrossed status unknown.

MT1079 *egl-15(n484)* X. Outcrossed once.

RB1091 *Y64G10A*.*7(ok1062)* IV. Not outcrossed.

LT121 *dbl-1(wk70)* V. Outcross status unknown.

KR827 *let-363(h502) dpy-5(e61) unc-13(e450)* I; *sDp2 (I;f)*. Outcross status unknown.

Additionally, the *E. coli* strain OP50-GFP was ordered from the *Caenorhabditis* Genetics Center (see Worm Maintenance).

### Worm maintenance

Worms were grown on solid agar nematode growth media (NGM) plates with ampicillin at 20°C. Temperature sensitive mutants were maintained at 15°C. Instead of OP50 *E. coli*, OP50-GFP, a strain of OP50 that contains a GFP plasmid (pFPV25.1) was used since it was resistant to ampicillin and allowed addition of the antibiotic in NGM plates (100 μg/ml). This measure was taken to minimize contamination as students began to learn how to maintain and experiment with *C. elegans*. As a weekly practice, students picked single worms onto seeded NGM plates and worked with the resulting population the following week.

### Orthology

To assess orthology between a *C. elegans* gene and the human proteome we use the DIOPT search engine https://www.flyrnai.org/cgi-bin/DRSC_orthologs.pl, which aggregates results from 16 different orthology matching programs. Orthology for the purpose of phenolog statistical analysis was based on a different method (see below).

### Phenolog statistical analysis

Human orthologs as well as RNAi and allele phenotypes associated with *C. elegans* genes were retrieved from WormBase https://wormbase.org/tools/mine/simplemine.cgi. Human phenotypes associated with human genes were retrieved from The Human Phenotype Ontology (HPO) project https://hpo.jax.org/data/annotations. Hypergeometric testing was performed in R (see Data and Code Availability).

### Tap response assay (areflexia)

Individual wildtype (N2) and *ndrr-2(ok1899)* (RB1565) worms were placed on unseeded NGM plates and allowed to acclimate for 2 minutes. A sterile eyelash attached to a pipette tip was used to deliver head taps. Each worm received 10 taps, with 15–20 seconds between each tap. The number of tail bends during withdrawal, omega turns and pauses were recorded and used to quantify reflex. *ndrr-2(ok1899)* (RB1565) was not outcrossed.

### Locomotion assay (hip flexor weakness)

Young adult *C. elegans* individuals maintained on seeded NGM plates were transferred to unseeded plates. For each worm, the number of complete body bends (forward or backward) was recorded over a 7-minute period. A two-sample *t*-test was used to compare body bend frequency between wildtype and *cca-1* mutants.

### Bursting assay (arterial rupture)

N2 and *let-2(ts)* larvae were maintained at 15°C, 20°C, or 25°C and then acclimated to room temperature the night before testing. Adult worms (5–7 days old) were transferred to individual wells of a 96-well plate containing 300 µL of distilled water. Worms were monitored under a microscope at 2.5-minute intervals for 10 minutes. Phenotypic categories included mobile, immobile, rod-like, or ruptured. The number of ruptured worms at each time interval was recorded for each condition.

### Identifying optimal temperature shift conditions (abnormal upper limb bone morphology)

Eight *let-2(ts)* worms were individually plated and shifted to 25°C at varying timepoints to determine the optimal temperature shift for survival. Plates shifted after 24 hours produced the highest number of viable adults.

### Viability and pharyngeal development (cleft soft palate)

Synchronized populations of wildtype N2 and *dbl-1(wk70)* worms were transferred to seeded NGM plates and incubated at 25°C. Larvae were counted daily for two days and dead embryos scored. Pharyngeal morphology was examined under a compound microscope (40 × magnification). A Welch two sample *t*-test was used to test the difference between wildtype and *dbl-1* since variance between the two samples was assumed to be different..

### Swimming Behavior Assay (abnormal cellular phenotype)

Synchronized young adult wildtype and *polg-1(ok1548)* heterozygous worms were grown on seeded NGM plates. Worms were transferred into droplets of M9 buffer on microscope slides. Swimming durations were recorded for each worm, with swimming defined as thrashing within a 30 second time window (i.e., if worms stopped moving for more than 30 seconds it was deemed no longer swimming).

### Embryo Staging by Microscopy (premature birth)

Adult worms were transferred to slides with deionized water or M9 buffer and ruptured using coverslip pressure or osmotic stress. Released embryos were categorized by cell stage. In some cases, embryos were counted while still inside the worm. Two worms with 3 embryos in their uteri each were considered outliers and excluded from downstream analysis.

### Digestive tract integrity (lactose intolerance)

Worms were washed off plates using M9 buffer and divided into three treatment conditions: control, 50 mmol/L lactose, and 100 mmol/L lactose. Each group was washed three times using centrifugation at 1000 rpm for 3 minutes. After washing, worms were stained with Neutral Red (75 mmol) in 20% acetone and 80% DI water, incubated in the dark for 1 hour, and then washed with DI water. Neutral Red has an excitation peak at approximately 530–540 nm and emits at 640–650 nm. The available fluorescence microscope was equipped with DAPI (excitation 350–370 nm, emission 420–480 nm), GFP (excitation 460–490 nm, emission 510–550 nm), and

TXR (excitation 540–580 nm, emission 600–660 nm) filter sets. Although none of these filter sets matched Neutral Red’s spectral properties exactly, the TXR filter provided the closest overlap with both its excitation and emission spectra. Thus, the TXR filter was adopted for imaging, enabling qualitative visualization of dye retention in the gut despite suboptimal excitation efficiency. Imaging parameters were standardized for exposure (192.1), gain (4.4), and magnification (40 ×).

### Morphometric Analysis (lethal short-limb stature)

Wildtype N2 and *egl-15(n484)* mutant worms were maintained on seeded NGM plates and selected for imaging at the young adult stage. Using a mobile phone, images were taken through a calibrated binocular microscope with a stage micrometer eyepiece (90 units = 2 millimeters). Magnification was set to 5 × and the microscope was focused on young adult worms before acquisition. Images were converted to TIFF format and manually analyzed using Fiji (ImageJ) to determine body length and widths at the vulva and pharynx. We chose to measure worms on an NGM plate at 5 × rather than on an agarose pad at 40 × in order to increase the number of assayed individuals for better statistical analysis.

## Results

### Weak response to tapping as a model for areflexia

Areflexia (HP:0001284) is the absence of neurologic reflexes such as the knee-jerk reaction [3]. Among the genes with inferred associations, *NDRG1* was selected to attempt modeling areflexia in *C. elegans*. The NDRG family derives its name from “N-myc downstream regulated gene” [12]. *C. elegans* has a different yet orthologous MYC network in which two Max orthologs, MXL-1 and MXL-3 act as central dimerization partners for the Mad-like ortholog MDL-1 and a single Mlx ortholog, MXL-2, acts as central dimerization partner for Mondo-like protein MML-1 [13]. While a *bona fide C. elegans* ortholog of *NDRG1* could not be ascertained since no reciprocal best hits were found, genes Y48G10A.3 and ZK1073.1 are homologs of *NDRG1, NDRG2* and *NDRG3*, with Y48G10A.3 also being an ortholog of *NDRG3* [14]. Furthermore, ZK1073.1 is a target of MDL-1 supporting its identity as an N-myc downstream regulated gene [15]. Since Y48G10A.3 and ZK1073.1 are “NDRG family member related” we named these two genes *ndrr-1* and *ndrr-2*, respectively.

We reasoned that if *C. elegans* could serve as a model for areflexia, then perhaps *ndrr-2* loss-of-function would impede the nematode’s ability to react to its surroundings. Therefore, we hypothesized that the knockout *ndrr-2(ok1899)* would show a dampened response in a tap-withdrawal assay compared to wildtype animals (**Methods**). To test this hypothesis we recorded the number of tail bends and omega turns after a series of ten separate head taps with an eyelash (n = 10 trials of ten taps for each strain). Additionally, we counted the number of times a worm paused while withdrawing from the tap, an indication of a dampened response (n = 8 trials of ten taps for each strain). In support of our hypothesis, we found that the mean number of tail bends was significantly reduced in *ndrr-2(ok1899)* animals (2.1 vs 3.4 tail bends per tap, *p*-value = 4.1 × 10^-12^). Likewise, we observed a significant reduction in total number of omega turns performed by each assayed worm (1.7 vs 3.8 omega turns during 10 taps, *p*-value = 1.2 × 10^-5^). Moreover, *ndrr-2(ok1899)* animals paused significantly more times during their withdrawal compared to wildtype worms (1.25 vs 1.13 pauses during 10 taps, *p*-value = 0.01) (**Figure 1a**). Taken together, these findings suggest that *ndrr-2* loss-of-function can recapitulate aspects of areflexia, providing a functional model for investigating motor and sensory neuropathy associated with *NDRG1* mutations in humans [16].

**Figure 1.**
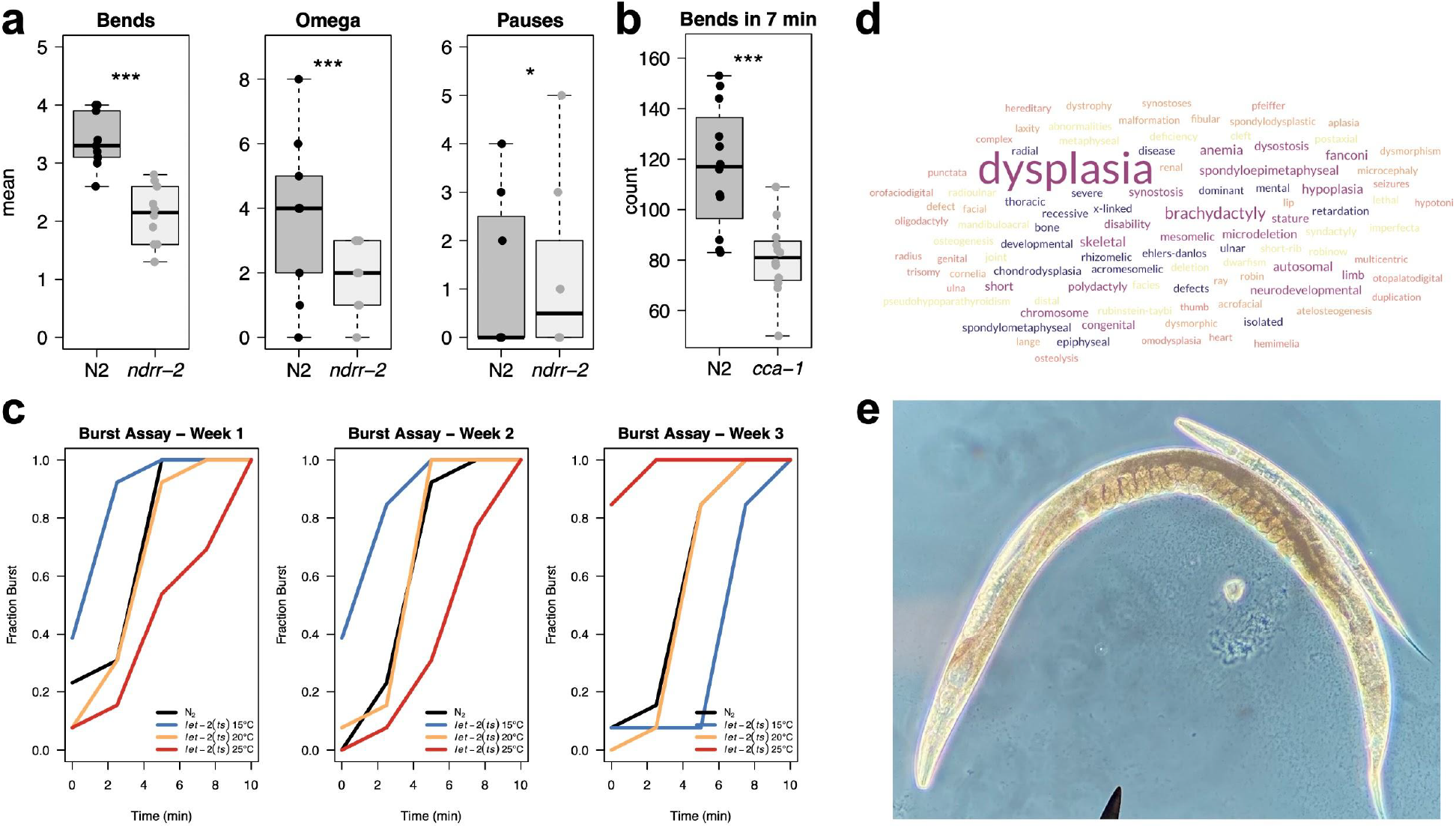
Modeling areflexia, hip flexor weakness, arterial stiffness and abnormal limb morphology in *C. elegans*. **a)** The mean number of tail bends and omega turns in wildtype and *ndrr-2(ok1899)* worms following ten evenly spaced head taps (n = 10 trials), and the mean number of pauses observed following ten evenly spaced head taps (n = 8 trials). Individual data points are overlaid on boxplots. Student’s *t*-test: *p* ≤ 0.01 (*); *p* ≤ 0.001 (***). **b)** The number body bends that wildtype and *cca-1(ad1650)* worms performed over a period of 7 minutes (n = 12 trials). Individual data points are overlaid on boxplots. Student’s *t*-test: *p* ≤ 0.001 (***). **c)** Burst assay results across three successive weekly trials using wildtype and *let-2(ts)* worms. In Weeks 1 and 2, worms reared at the permissive temperature (15°C) appeared to rupture more readily, likely due to uncontrolled temperature-dependent effects on cuticle elasticity. In Week 3, incorporation of a room temperature acclimation step demonstrated that *let-2(ts)* worms raised at the restrictive temperature (25°C) ruptured significantly faster than those raised at 15°C. **d)** A word cloud of the disease terms associated with Abnormal upper limb bone morphology. **e)** A wildtype (larger) and *let-2(ts)* worm 4 days after hatching displaying the stark difference in growth rate. The wildtype worm has reached adulthood as evidenced by the embryos developing in its uterus. The *let-2(ts)* worm was hatched and reared at permissive temperature (15°C) for 24 hours before being shifted to the restrictive temperature (25°C).

### Impaired sinusoidal movement as model for hip flexor weakness

Hip flexor weakness (HP:0012515) is the reduced ability to flex the femur, that is, to pull the knee upward [3]. Among the genes with inferred associations, *SCN4A* was selected to model hip flexor weakness in *C. elegans. SCN4A* encodes a voltage-gated sodium channel and is responsible for sodium ion conductance in muscle cells [12]. Unbeknownst to us when selecting *SCN4A*, the *C. elegans* genome does not encode any voltage-gated sodium channels [17]. Instead, voltage-gated calcium channels serve an equivalent role in translating electrical signals into muscle contractions. Therefore, to model hip flexor weakness as a result of cation channel loss-of-function the *C. elegans* voltage-gated calcium channel *cca-1* was adopted [18].

We drew an analogy between impaired gait in humans due to hip flexor weakness and impaired locomotion in nematodes due to reduced muscle activity. Consequently, we hypothesized that the number of sinusoidal wave motions will be reduced in *cca-1* mutants. To test this hypothesis we recorded the number of body bends that wildtype and *cca-1(ad1650)* worms perform over a period of 7 minutes (n = 12 trials each) and found that the mean number of bends was significantly reduced in *cca-1(ad1650)* (81 vs. 117 bends, *p*-value = 5.4 × 10^-15^) **(Figure 1b and Methods)**. This finding supports the hypothesized importance of *cca-1* in locomotion and suggests that its disruption can model neuromuscular phenotypes such as hip flexor weakness.

### Worm bursting as a model for arterial rupture

Arterial rupture (HP:0025019) is the sudden breakage of an artery leading to leakage of blood from the circulation [3]. Among the genes with inferred associations, *COL5A2* was selected to model arterial rupture in *C. elegans. COL5A2* encodes a collagen alpha-2(V) chain and contributes to extracellular matrix structure and stability [12]. In *C. elegans, let-2* encodes a structural collagen involved in maintaining cuticle integrity [19]. While not an ortholog of *COL5A2*, we chose *let-2* to model arterial rupture due to compromised collagen function.

We reasoned that reduced cuticle integrity in nematodes could lead to breakage reminiscent of arterial rupture in humans. Consequently, we hypothesized that the internal pressure resulting from placing *let-2(ts)* worms in a hypotonic solution after rearing them at the restrictive temperature would lead to cuticle breakage and worm bursting. Initial trials yielded inconsistent results, likely due to uncontrolled effects of temperature on tissue elasticity (**Figure 1c**). Therefore, we incorporated an acclimation step at room temperature before each trial (**Methods**). The revised assay revealed that *let-2(ts)* worms raised at the restrictive temperature (25°C) ruptured significantly faster than those raised at the permissive temperature (15°C), as indicated by a log-rank test (χ^2^(1) = 23.7, *p* = 1 × 10^-6^) (**Figure 1c**).

This finding indicates that *let-2(ts)* worms exhibit temperature-dependent sensitivity to osmotic pressure, consistent with defects in structural integrity and supports the use of *let-2(ts)* as a model for exploring arterial rupture due to compromised collagen function.

### Worm body size as a model for abnormal limb bone morphology

Abnormal upper limb bone morphology (HP:0040070) refers to the atypical shape or structure of the bones that make up the arm, which could impact function of the arm or hand [3]. According to The Human Phenotype Ontology (HPO) website, this phenotype is associated with over 800 human diseases. A word cloud of the disease terms highlights a common theme of dysplasia, a usually neoplastic transformation of the cell, associated with altered architectural tissue patterns **(Figure 1d)**. From among the genes associated with abnormal upper limb bone morphology, *COL10A1* was selected for further study. The protein encoded by *COL10A1* is 680 amino acids long alpha-1 chain of type X collagen that affects cartilage and endochondral osteogenesis [12,20].

Although nematodes do not develop limbs, we surmised that abnormal upper limb morphology could nevertheless be modeled in *C. elegans*. We reasoned that since abnormal limb bone morphology could represent an misexpression of the *COL10A1* gene during osteogenesis then misexpression of collagen in the worm could lead to an analogous phenotype during cuticle formation. *The C. elegans* ortholog of COL10A1 is *col-104*, however we adopted the homolog *let-2* to investigate the effects of collagen misexpression to take advantage of a temperature sensitive mutant [19]. We hypothesized that *let-2* mutants would display atypical shapes or structures in the cuticle when grown at a restrictive temperature.

We shifted a temperature sensitive *let-2* mutant to restrictive temperatures at different developmental stages to determine conditions that allowed survival to adulthood **(Methods)**. *let-2(ts)* worms shifted from 15°C to 25°C 24 hours after fertilization showed reduced lethality and were used to score for abrupt angulations in the worm body. We noted that one-day-old *let-2(ts)* larvae shifted to, and maintained at, restrictive temperature until adulthood were significantly shorter in body length compared to wildtype controls **(Figure 1e)**. Due to time constraints, this phenotype was not thoroughly quantified. However, the observation of short *let-2* worms led us to explore performing a phenolog analysis [21]. We asked if a significant number of orthologs associated with abnormal upper limb bone morphology in humans were also associated with the short phenotype in *C. elegans*. Hypergeometric testing revealed that indeed the overlap in orthologs associated with these two phenotypes was statistically significant (*N* = 16,329, *n* = 375, m = 145, c = 22, *p*-value = 2.1 × 10^-12^). Together, these results suggest that reduced body length in *let-2* mutants can serve as a proxy for modeling limb morphologies associated with collagen defects in humans.

### Modeling cleft soft palate via pharyngeal development in *C. elegans*

Cleft soft palate (HP:0000185) is a result of a developmental defect that leads to malformation of soft tissue structures in the oral cavity [3]. Among the genes with inferred associations, *TGFB3* was selected to model cleft soft palate in *C. elegans*. Although *dbl-1* is not a *bona fide* ortholog of *TGFB3* it is the best alignment match and shares functional similarity as ligands in TGF-β signaling pathway [12,14].

We hypothesized that *dbl-1* mutants would display defective pharyngeal development and morphology reminiscent of a cleft soft palate. To test this hypothesis, the offspring of individual N2 and *dbl-1(wk70)* animals were scored for embryonic lethality (N = 3 parents for each genotype). The mean number of offspring from *dbl-1(wk70)* moms was significantly lower than wildtype (41±31.2 vs. 161±11.4 animals, t = 5.9312, df = 2.4631, *p*-value = 0.016). However, no embryonic lethality nor signs suggestive of pharyngeal maldevelopment in larvae were observed at 25°C (**Methods**). Consistent with published findings, *dbl-1(wk70)* animals were reduced in size underscoring the importance of this gene for growth and development [22]. Although a pharynx-specific phenotype was not detected, the reduced growth observed in *dbl-1(wk70)* encourages further investigation into whether this gene can be used to model aspects of soft tissue developmental defects associated with *TGFB3* dysfunction.

### Modeling mitochondrial dysfunction through impaired motility in *C. elegans*

Abnormal cellular phenotype (HP:0025354) ecompasses a wide range of anomalies of cellular morphology or physiology [3]. Among the hundreds of genes associated with this phenotype we chose to study *POLG*, which encodes the mitochondrial DNA polymerase [12]. In *C. elegans*, the ortholog *polg-1*, was adopted for modeling abnormal cellular physiology caused by mitochondrial dysfunction [23].

We reasoned that *polg-1* mutants would suffer from energy deficits due to disrupted mitochondrial function. Indeed, worms homozygous for a *polg-1(ok1548)* mutation are sterile and have shortened lifespan [24]. Therefore, we hypothesized that *polg-1* heterozygous animals would have impaired mitochondrial activity leading to reduced physical fitness in the worm. To test this hypothesis, we conducted assays to measure how long worms could swim **(Methods)**. Surprisingly, *polg-1(ok1548)* heterozygotes swam for significantly longer periods than wildtype worms! Worms with one copy of mutated *polg-1* swam on average for 6.4 minutes compared to 4.9 minutes swam by N2 individuals **(Figure 2a)**. A Wilcoxon rank-sum test indicated that the observed difference was significant (Wilcoxon rank-sum test, W = 564, *p*-value = 2.3 × 10^-6^). Importantly, this result was confirmed independently by two students (Wilcoxon rank-sum test, W = 147, p = 0.019 and W = 133, p = 0.00032) thereby adding credence to the counterintuitive enhanced swimming fitness in *polg-1* heterozygotes. These results rule out the hypothesized consequence of *polg-1* loss-of-function and is interpreted as a form of systemic overcompensation for mitochondrial deficiency.

**Figure 2.**
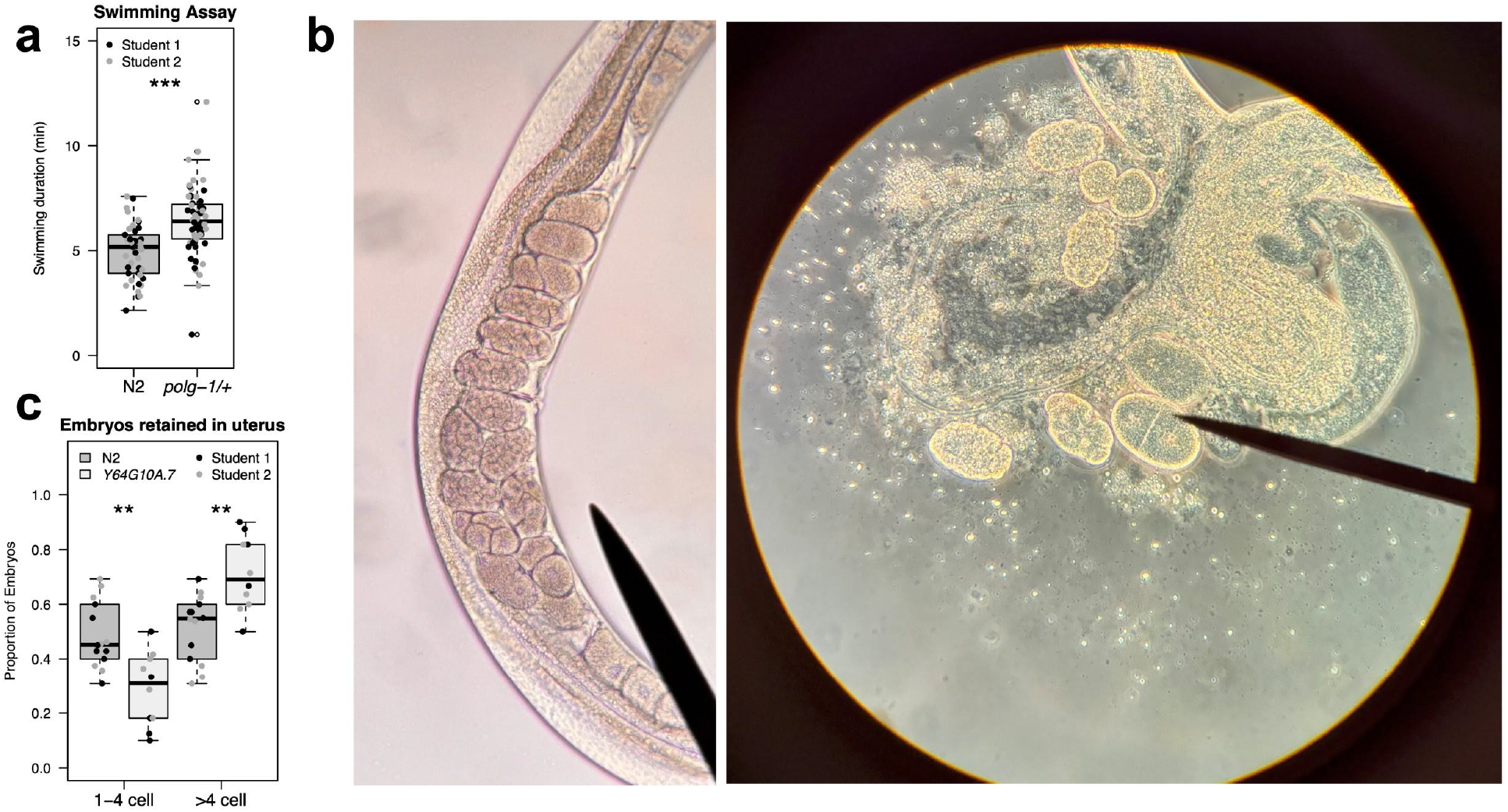
Modeling abnormal cell morphology and premature birth in *C. elegans*. **a)** Swimming durations of wildtype and *polg-1(ok1548)*/+ heterozygous worms. Boxplots represent 42 wildtype and 60 *polg-1* heterozygous individuals. Data points reflect two durations measured by two different researchers, both of which independently confirmed the enhanced swimming phenotype in *polg-1* heterozygotes. Statistical significance was assessed using the Wilcoxon rank-sum test: *p* ≤ 0.001 (***). **b)** An example of the setup used to quantify egg retention. Embryonic stages were quantified *in utero* or *ex utero* after worm rupture. The needle in the image to the right points to a two-cell stage embryo. **c)** The proportions of embryos with few (i.e. 1 to 4 cells) versus many (i.e., > 4 cells) in 28 wildtype and 20 *Y64G10A*.*7(ok1062)* young adults. Data points reflect values ascertained by two independent researchers. Wilcoxon rank-sum test: *p* ≤ 0.01 (**).

### Egg retention dynamics as a model for premature birth

Premature birth (HP:0001622) is the birth that occurs before 37 weeks of gestational age [3]. Among the genes with inferred associations, *LTBP3* was chosen to attempt and model human premature birth in *C. elegans*. LTBP3 encodes Latent Transforming Growth Factor Beta Binding Protein 3, a protein rich in EGF-like domains and a β-sheet core and is a regulator of TGF-beta signalling [12]. *LTBP3* does not have any orthologs in *C. elegans* [14]. However, the top match is the uncharacterized *Y64G10A*.*7* (26.8% identity, E-value = 1.3 × 10^-61^), which in turn is a *bona fide* ortholog of the human gene MEGF6 (Multiple EGF Like Domains 6) [14].

Despite the fact that *C. elegans* is an oviparous species that lays its embryos rather than retain and nourish them *in utero*, we wondered if egg retention versus premature egg laying could be an analogous phenotype to premature birth in humans. We hypothesized that *Y64G10A*.*7* loss-of-function would lead to premature egg-laying, detectable by an increased number of early-stage embryos retained in the uterus. To test this, gravid worms from both wildtype N2 and the knockout strain *Y64G10A*.*7(ok1062)* were ruptured between an agarose pad and coverslip and the developmental stages of released embryos were categorized based on cell number (**Methods and Figure 2b**). Contrary to our hypothesis, wildtype *C. elegans* retained a significantly higher proportion of early-stage embryos (1-cell to 4-cell stage) compared to the *Y64G10A*.*7(ok1062)* mutant (Wilcoxon rank-sum test, W = 120.5, p = 0.003; **Figure 2c**). This finding suggests that loss of *Y64G10A*.*7* function may promote retention of later-stage embryos, potentially acting as a negative regulator of egg-laying. Consequently, this result questions whether the inferred association between human *LTBP3* and premature birth reflects a positive or negative association and encourages further exploration of TGF-β signaling in developmental timing.

### Modeling lethal short-limbed short stature *C. elegans* morphometrics

Lethal short-limbed short stature (HP:0008909) is also known as lethal micromelic dwarfism [3]. The only gene hitherto associated with this phenotype is Fibroblast Growth Factor Receptor 3 (*FGFR3*), which encodes a transmembrane tyrosine kinase receptor that plays a critical role in chondrocyte proliferation and skeletal development [3,12]. The *C. elegans* ortholog of FGFR3 is *egl-15*, which also encodes a fibroblast growth factor receptor critical for development and cell migration [25]. Therefore, we aimed to model lethal micromelic dwarfism in *C. elegans* by studying the consequences of *egl-15* loss-of-function.

We hypothesized that *egl-15(n484)* would result in reduced body length and altered body shape, recapitulating aspects of short-limbed dwarfism. To test this, young adult worms were imaged using a calibrated stereo microscope **(Methods and Figure 3a)**. Morphometric data were collected for total body length and for body widths at two anatomical landmarks: the pharynx and the vulva. We found no significant difference between the mean lengths of wildtype and mutant animals (0.98±0.11 vs 0.94±0.15 mm, *p*-value = 0.39, Wilcoxon rank sum test) or between the width at the pharynx (35.9±7 vs 39.6±11 microns, *p*-value = 0.13), although a modestly significant difference in body width at the vulva was detected (55.7±9 vs 60.4±17 microns, *p*-value = 0.043). However, when width at pharynx or vulva were normalized to the length of an individual a more compelling difference was detected in both cases (pharynx: 35.3±6 vs 41.7±11 microns, *p*-value = 0.022 and vulva: 56.4±5 vs 63.2±12 microns, *p*-value = 0.0008958) (**Figure 3b**). Therefore, while there was no significant difference in absolute body length, overall shape was different between wildtype and mutants with *egl-15(n484)* being relatively broader.

**Figure 3.**
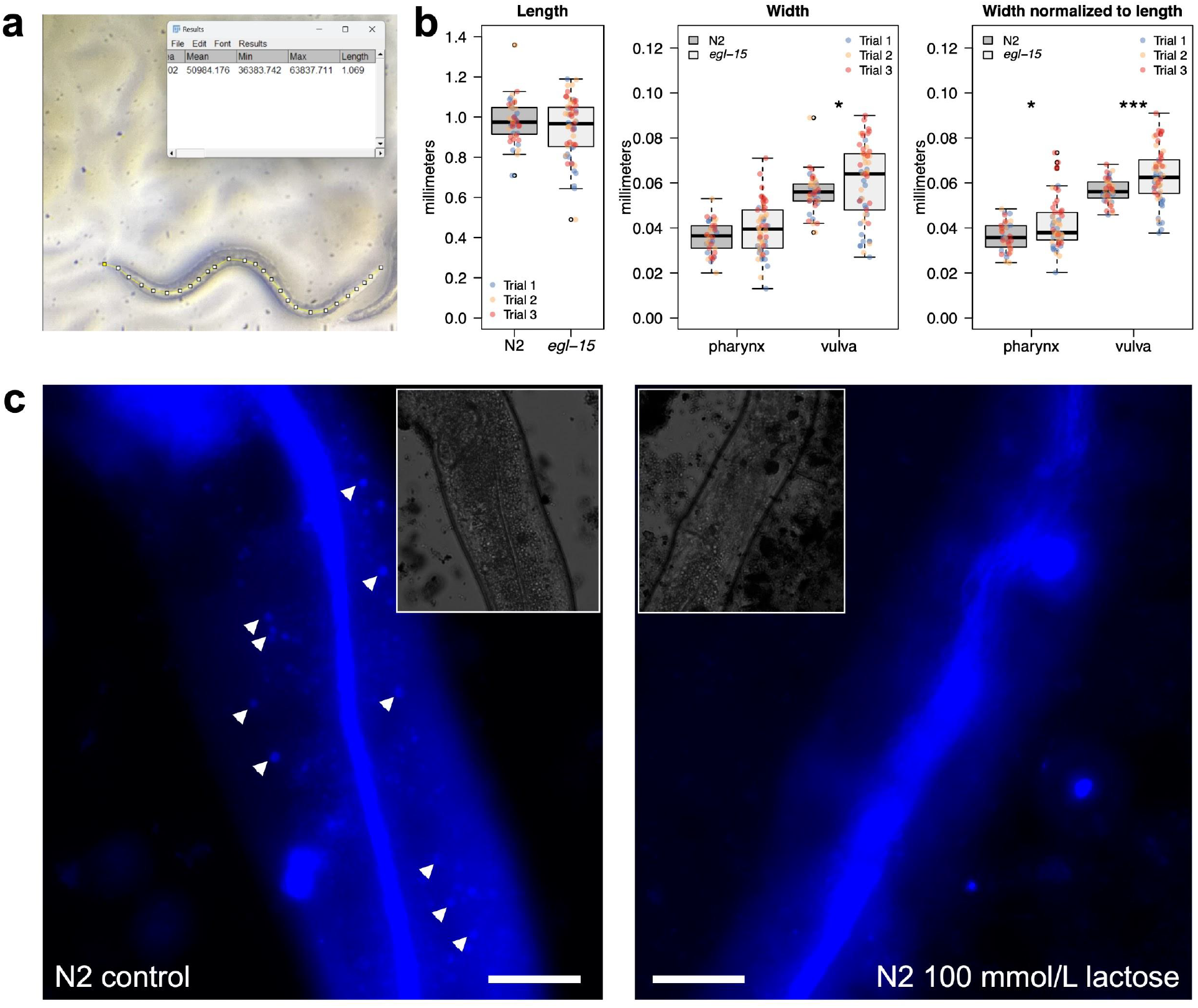
Modeling lethal short-limb status and lactose intolerance in *C. elegans*. **a)** An example of the interface used for *in situ* quantification of worm morphometrics in Fiji. **b)** Morphometric data for wildtype and *egl-15(n484)* animals across three successive weekly trials were collected for total body length and for body widths at pharynx and vulva. Length was measured in 44 wildtype and 59 *egl-15* worms; pharynx width in 38 and 58; and vulva width in 40 and 58, respectively. Wilcoxon rank sum test: *p* ≤ 0.01 (*); *p* ≤ 0.001 (***). **c)** Representative fluorescence microscopy images of wild-type *C. elegans* intestine stained with neutral red. Brightfield images shown in insets. Arrowheads indicate visible gut granules.

These findings are consistent with the known role of *FGFR3* in regulating tissue elongation and skeletal growth and highlights the potential of *C. elegans* to capture key aspects of growth impairment and body proportion changes associated with skeletal dysplasias.

### Gut integrity as model for lactose intolerance

Lactose intolerance is a human metabolic phenotype (HP:0004789) characterized by the inability to digest lactose [3]. Among the genes with inferred associations, *mTOR* was selected to model lactose intolerance in *C. elegans*. The ortholog of *mTOR* in *C. elegans* is *let-363* and similar to mTOR, the encoded protein is a member of the PI3/PI4K kinase family and shares a similar domain architecture, supporting functional conservation and justifying its selection to model lactose intolerance in *C. elegans* [26].

We hypothesized that loss of *let-363* function would reduce gut integrity in *C. elegans* in response to a lactose rich diet. To test this, we first adapted protocols to treat both wildtype and *let-363* mutant worms with lactose and then asses gut integrity by staining with neutral red, a vital dye that accumulates in lysosome-rich gut cells and serves as a marker of tissue viability [27]. Using the developed protocol we could detect a difference in the number of fluorescent gut granules when comparing untreated N2 worms to worms treated with 100 mmol/L lactose (**Methods and Figure 3c**). Due to time constraints, we could not quantify the difference in observed gut granules, determine the contribution of autofluorescence or compare with a *let-363(h502)* mutant. Nevertheless, the developed protocol provides a starting point for future students to pursue this experiment and determine the plausibility of using *let-363* as a model for lactose intolerance resulting from mTOR loss-of-function.

## Discussion

This study presents the scientific output of a research-based undergraduate learning module. Through nine independent experiments, students explored the conservation of molecular and phenotypic traits and gained hands-on experience in experimental design and data interpretation. A key significance of this work lies in empowering undergraduates to engage as active contributors to scientific inquiry and to share their contributions with the broader scientific community.

The use of *C. elegans* proved highly effective for undergraduate education due to its genetic accessibility and well-documented developmental biology. The challenge of modeling aspects of complex human development in a nematode prompted students to trace the common ancestry of seemingly distant phenotypes. Such “homology chasing” offered opportunities to see beyond visible phenotypes and deeper into layers of conserved and interconnected genetic effects. Additionally, developing and troubleshooting protocols within the allotted time period required students to cooperate, multitask and practice prudent time management. Weekly evaluation of worm colonies in a transparent manner using a common spreadsheet aimed to instill an openness with regards to success and failure (e.g., starved worms) and a willingness to mitigate shortcomings (e.g., better worm maintenance). Engagement with WormBase, AlphaFold, Uniprot and the Human Phenotype Ontology database further equipped students to navigate and use increasingly important resources. Invariably, each group experienced the satisfaction of overcoming an obstacle or the surprise of an unexpected experimental outcome. Such a sense of satisfaction and associating it with methodical and open scientific inquiry is one of the key goals this educational module aimed to achieve.

By compiling these experiments into a publishable manuscript, this work challenges the traditional ephemerality of undergraduate research outputs and positions students as emerging scientists whose work merits global dissemination. This approach not only validates their contributions but also integrates undergraduate research into the broader scientific endeavor.

The rapid delivery of strains from the *Caenorhabditis* Genetics Center and guidance from WormBase on gene nomenclature were critical to the project’s success, highlighting the importance of community resources in supporting both research and education.

Limitations, such as time constraints and suboptimal assay conditions (e.g., neutral red imaging using Texas Red filter), underscore areas for future refinement. Nevertheless, these challenges provided valuable lessons in resourcefulness and adaptability. This work advocates for a model of undergraduate science education where students are active contributors driven by curiosity and empowered to share their findings with the world. By demonstrating the feasibility and impact of such an approach, this study lays the foundation for future curricula that inspire and equip the next generation of scientists.

## Data and Code Availability

Data and the R script used for statistical analysis and figure generation is available at https://github.com/elewahub/homo_elegans

## Acknowledgements

We thank Matthew Beckman and Chris Palahniuk for support throughout the semester. The *Caenorhabditis* Genetics Center offered strains in a timely manner thereby enabling the success of this educational module. WormBase (Tim Schedl and Stavros Diamantakis) guided the process of naming ZK1073.1 and Y48G10A.3 and provided an intuitive platform for students to learn about their genes of interest. David Greenstein (University of Minnesota) recommended using twist ties for worm picking which proved to be a suitable and economical alternative to platinum wire picks.

## Author Contributions

AE taught the developmental biology course and instructed students on how to work with *C. elgeans*. EJ and LM investigated weak response to tapping as a model for areflexia. DK and YD investigated worm bursting as a model for arterial rupture. LY investigated impaired sinusoidal movement as a model for hip flexor weakness. AS1, AH and SM investigated abnormal worm body size as a model for human limb malformation and AE performed the phenolog statistical analysis. MC investigated modeling cleft soft palate via pharyngeal development in *C. elegans*. AS2 and TW investigated modeling mitochondrial dysfunction through impaired motility in *C. elegans*. AV and PX investigated egg retention dynamics as a model for premature birth. BL and DS investigated modeling lethal short-limbed short stature via morphometrics in *C. elegans*. MS and DW investigated modeling lactose intolerance. All authors participated in writing the manuscript together with AE.

